# Identification of novel candidate genes associated with the symbiotic compatibility of soybean with rhizobia under natural conditions

**DOI:** 10.1101/2024.04.04.587839

**Authors:** Masayoshi Teraishi, Kosuke Sakaguchi, Takahiro Tsuchimoto, Takanori Yoshikawa

## Abstract

A robust symbiotic relationship between soybean and rhizobia can enhance the yield and quality of soybeans by reducing nitrogen fertilizer inputs, thereby contributing to sustainable agriculture. The genetic interplay between soybean cultivars and the rhizobial species colonizing their roots under natural conditions remains underexplored. This study builds on the observation that the prevalence of rhizobial species associated with the soybean cultivars ‘Peking’ and ‘Tamahomare’ varies significantly. Herein, we performed a quantitative trait loci (QTL) analysis of the proportion of *Rhizobium* species present in the root nodules of these cultivars using recombinant inbred lines derived from a cross between ‘Peking’ and ‘Tamahomare.’ A major QTL was identified on chromosome 18, accounting for 42% of the phenotypic variation, and was subsequently localized to a 240 kb region. The RNA-seq analysis indicated that a single gene featuring nucleotide binding site–leucine-rich repeat domains exhibited markedly different expression levels in parent cultivars within the QTL region. As this locus is distinct from the chromosomal regions containing known nodule-related genes, such as *Rj* or *rj*, it likely represents a novel gene involved in symbiosis between rhizobia and soybeans. Further research of the function and role of this new gene has potential to improve soybean yield and contribute to sustainable agriculture under low nitrogen fertilizer conditions.

## 1 INTRODUCTION

Soybean (*Glycine max*) is one of the most economically important crops worldwide. It is highly suitable for human and animal diets and is a source of 30% of the world’s oil derived from processed crops. Soybean originated in northern China and is expanding worldwide, particularly in North and South America. The use of soybean in biofuel production is further increasing its economic impact. The rising economic importance of soybean has led to increased efforts over the past several years to improve soybean productivity.

Legume plants, including soybeans, exhibit the unique characteristic of symbiosis with rhizobia for nitrogen fixation (Doyle & Luckow, 2003). Nitrogen fixed by legumes is a major nitrogen source in soil (Schaedel et al., 2021). Legume crops occupy 12–15% of the arable land worldwide (Graham & Vance, 2003), and nitrogen fixation in legume crops provides 40 million tons of nitrogen annually to agricultural land (Galloway et al., 2008). The N-fixing capacity of legumes contributes to the nitrogen supply of both natural and arable ecosystems. The effective utilization of nitrogen-fixing capacity is necessary for developing sustainable and low-input agriculture.

Several genetic factors, called *Rj* or *rj* genes, which regulate nodulation traits, have been identified in soybeans. They were identified under inoculation conditions using specific rhizobial strains. Genetic factors under uninoculated and natural conditions have not been investigated effectively.

Our previous research (Ramongolalaina et al., 2018), performed in a natural environment under non-inoculated conditions, showed that novel genes located on chromosome 18 in soybeans control the compatibility of rhizobial strains with host plants. No genes responsible for nodulation were identified in this region. Here, we narrowed down the quantitative trait loci (QTL) region and selected candidate genes using RNA-seq analysis.

## 2 MATERIALS AND METHODS

### 2.1 Plant materials

In this study, we used 113 F_6_-derived RILs, derived from a cross between their parent cultivars, ‘Tamahomare’ and ‘Peking.’ The major nodulating rhizobial species in the root nodules of ‘Tamahomare’ was *B. japonicum*, whereas the dominant rhizobial species in the root nodules of ‘Peking’ was *B. elkanii*.

Soybean seeds of each line, sterilized by soaking in 70% ethanol for 30 s and 2.5% sodium hypochlorite solution for 3 min, were sowed in a pot (10 cm x 20 cm x 15 cm) filled with soil derived from soybean planted field in a Kyoto University experimental farm and grown in an incubator set at 28□ with 18/6 h photoperiod. Plants were removed from the soil at the fourth trifoliolate stage, a month after sowing. Twenty-four nodules per line were randomly harvested from roots.

### 2.2 DNA extraction

The nodules were washed to remove the soil, surface-sterilized with 70% ethanol for 1 min and 2.5% sodium hypochlorite solution for 3 min, and rinsed with sterilized water. Subsequently, the nodules were put into tubes one by one, submerged in 300 µl of TPS buffer containing 100 mM Tris-HCl, 1 M KCl, and 10.0 mM EDTA, and then crushed with multi-beads shocker (Yasui Kikai Co., Japan). After centrifugation at 10000 g for 1 min, the DNA in the supernatant was treated with isopropanol, washed twice with 70% ethanol, and dissolved in TE buffer.

### 2.3 PCR-based amplification and the classification of rhizobia species

The DNA sequence analysis of the ITS region between the 16S and 23S ribosomal RNA (rRNA) genes of nodule bacteria was used to classify the rhizobial species. In a previous report (Ramongolalaina et al., 2018), the classification of rhizobia species was based on restriction fragment length polymorphisms of the PCR products of the ITS region between the 16S and 23S rRNA genes. In this study, however, we adopted Illumina MiSeq-based high-throughput analysis using the sequence information of a part of the ITS region for classifying nodule bacteria.

A conserved tRNA-Ala sequence within the ITS region of *Bradyrhizobium* spp. was used for designing the forward primer for the first round of PCR, whereas the reverse primer was designed using the ITS-R sequence at the end of the 23S rRNA gene (Akao, 2004). An eight-base index, i.e., the i5/i7 index, was added between the first primer and P5/P7 sequence (Figure S5). Primers used for amplicon sequencing are listed in Table S4. Twenty-four forward primers and 24 reverse primers were mixed to create 576 primer sets containing 10.0 µM concentration of each primer.

For the first PCR analysis, 0.1 µL of DNA template was mixed with 10 µL of EmeraldAmp MAX PCR Master Mix (TaKaRa Bío. Co., Japan) and 0.2 µL of a primer set containing 10.0 µM concentration of each primer, and the volume was made up to 20 µl with distilled water. The PCR cycle consisted of a pre-run at 96□ for 2 min, denaturation at 98□ for 20 s, annealing at 62□ for 15 s, and extension at 72□ for 1 min. The sequence was repeated for a total of 25 cycles and followed by a final post-run for 1 min at 72□. The PCR products were purified using the LaboPass Gel Extraction Kit (Hokkaido System Science Co., Japan) after electrophoresis and mixed with 576 PCR products in one tube, followed by amplicon sequencing on the MiSeq platform (2 × 300 bp) (Illumina, USA) conducted by Bioengineering Laboratory Co. Ltd. (Japan).

### 2.4 Estimation of rhizobial species

Primer and index sequences were trimmed from fastq files using Cutadapt (Martin, 2011), quality filtering was performed using fastp (Chen et al., 2018), and the processed fastq files were divided into RILs. The manifest file was imported into QIIME2 (Bolyen et al., 2019). Classification of nodule bacteria was performed using QIIME2 based on known *Bradyrhizobium* ITS sequences.

### 2.5 Construction of linkage map

A linkage map of RILs was constructed using the GRAS-Di technology developed by TOYOTA (Enoki & Takeuchi, 2018). The DNA samples of RILs were sent to Eurofins Genomics (Japan), which performed genotyping using GRAS-Di technology. Co-dominant DNA markers accrued by GRAS-Di were used for linkage map construction using MapMaker 3.0 (Lander et al., 1987).

The QTL analysis was performed using the composite interval mapping function of the R/qtl package and the Haley–Knott regression method. The LOD significance threshold for detecting QTLs was calculated by performing 1,000 iterations using the R/qtl permutation test.

### 2.6 RNA-seq analysis

For RNA-seq analysis, total RNA was extracted with RNAiso Plus (TaKaRa Bio Co., Japan) from soybean roots cultivated on soil for a month and purified with RNeasy Mini Spin Columns (Qiagen, Japan) following the manufacturer’s protocols. Purified RNA was sent to the Bioengineering Lab. Co. Ltd. (Japan). The RNA-seq data were analyzed using the RaNA-Seq program (https://doi.org/10.1093/bioinformatics/btz854).

## 3 RESULTS

### 3.1 Analysis of QTL

Ramongolalaina et al. (2018) reported a major QTL associated with the compatibility of indigenous rhizobia on chromosome 18 using 93 recombinant inbred lines (F_14_-derived) (RILs) derived from a cross between ‘Peking’ and ‘Tamahomare.’

We redeveloped another set of RILs, consisting of 113 F_6_-derived between ‘Peking’ and ‘Tamahomare,’ and subjected them to high-resolution linkage mapping using genotyping by random amplicon sequencing-direct (GRAS-Di) technology. The linkage map had a total length of 2,846.3 cm and contained 860 markers, with, and the average distance between the markers was 3.3 cm (Table S1).

Rhizobial strains attached to the roots were divided into two groups, i.e., *Bradyrhizobium japonicum* and *Bradyrhizobium elkanii*, based on the amplicon sequence analysis of the partial internal transcribed spacer (ITS) region between tRNA-Ala and the end of 23S rRNA. The results of the QTL analysis in terms of the ratio of rhizobial strains showed that a single major QTL associated with the compatibility of indigenous rhizobia in soil was located between A08389 (52,593 kbp) and A35793 (57,317 kbp) on chromosome 18, responsible for 42% of the phenotypic variation (Table 1; Figure S1). The frequency distribution of RILs for the ratio of rhizobial strains between the two homozygous genotype groups in the QTL region showed a clear bias (Figure S2). The ‘Peking’ homozygous RILs showed tended toward a high frequency of *B. elkanii*, whereas the ‘Tamahomare’ homozygous RILs showed a high frequency of *B. japonicum*. The highest limit of detection (LOD) value was observed between A01978 (56,531 kbp) and A35793 (57,317 kbp) (Figure S3). A previous study (Ramongolalaina et al., 2018) detected QTLs in the same region near the simple sequence repeat marker Sat_064 (56,334 kbp) on chromosome 18.

**Table 1.**
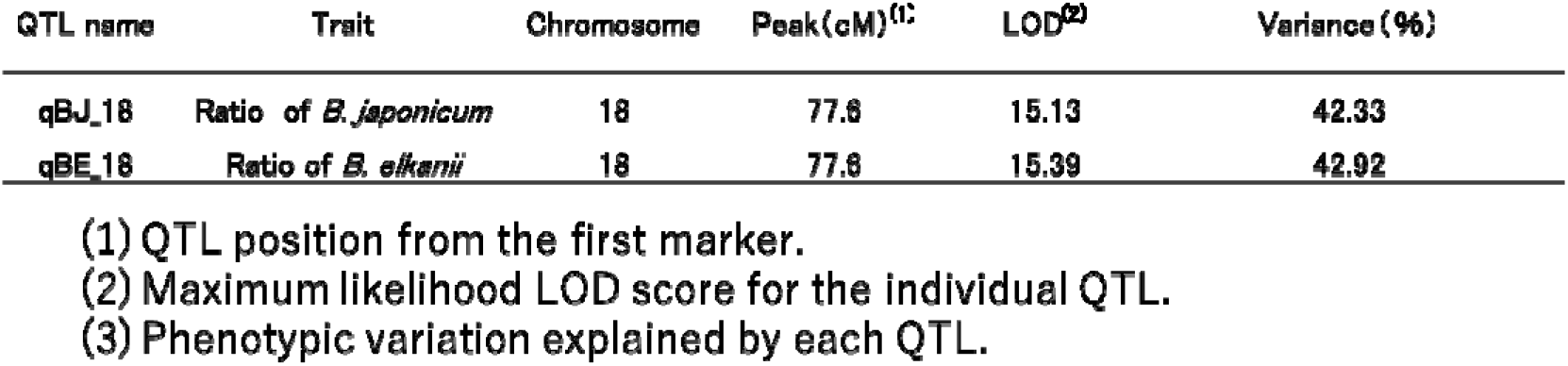
QTLs detected for nodule number ratios of *B. japonicum* and *B. elkanii*.

### 3.2 Fine mapping of the QTL

The RILs harboring recombination within the QTL region were selected, and additional polymerase chain reaction (PCR)-based markers for detailed genotyping were designed (Table S2) to narrow down the QTL.

Finally, the location of genes determining compatibility with rhizobia was narrowed from 56,433 kbp to 56,675 kbp on chromosome 18 (Figure 1). There were 22 predicted gene models, named Glyma.18g280800-Glyma.18g283100, based on the genome assembly sequence data of Williams 82 (Glyma.Wm82.a2) (Table 2).

**Table 2.**
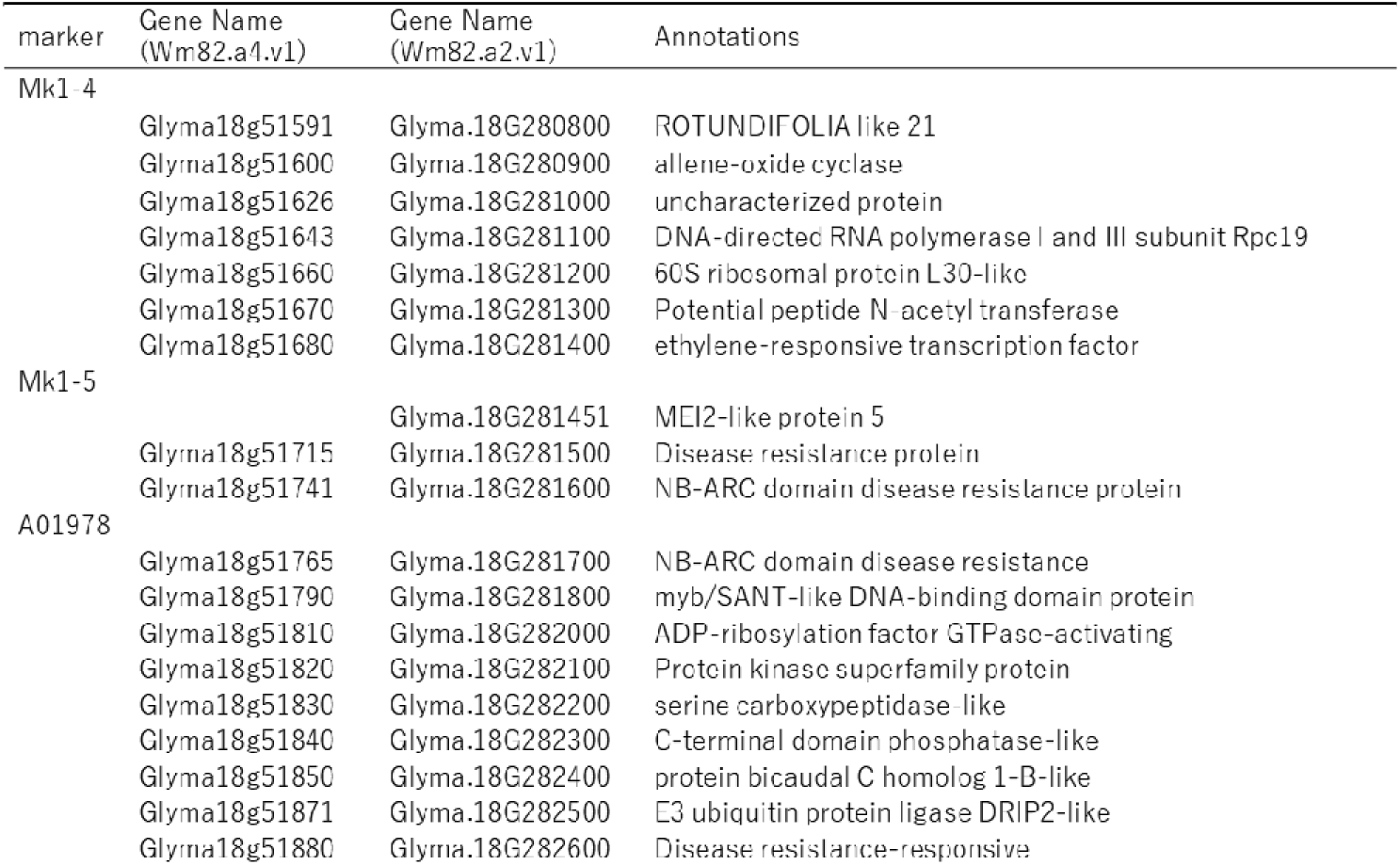
Candidate genes within the narrowed QTL region.

**FIGURE 1.**
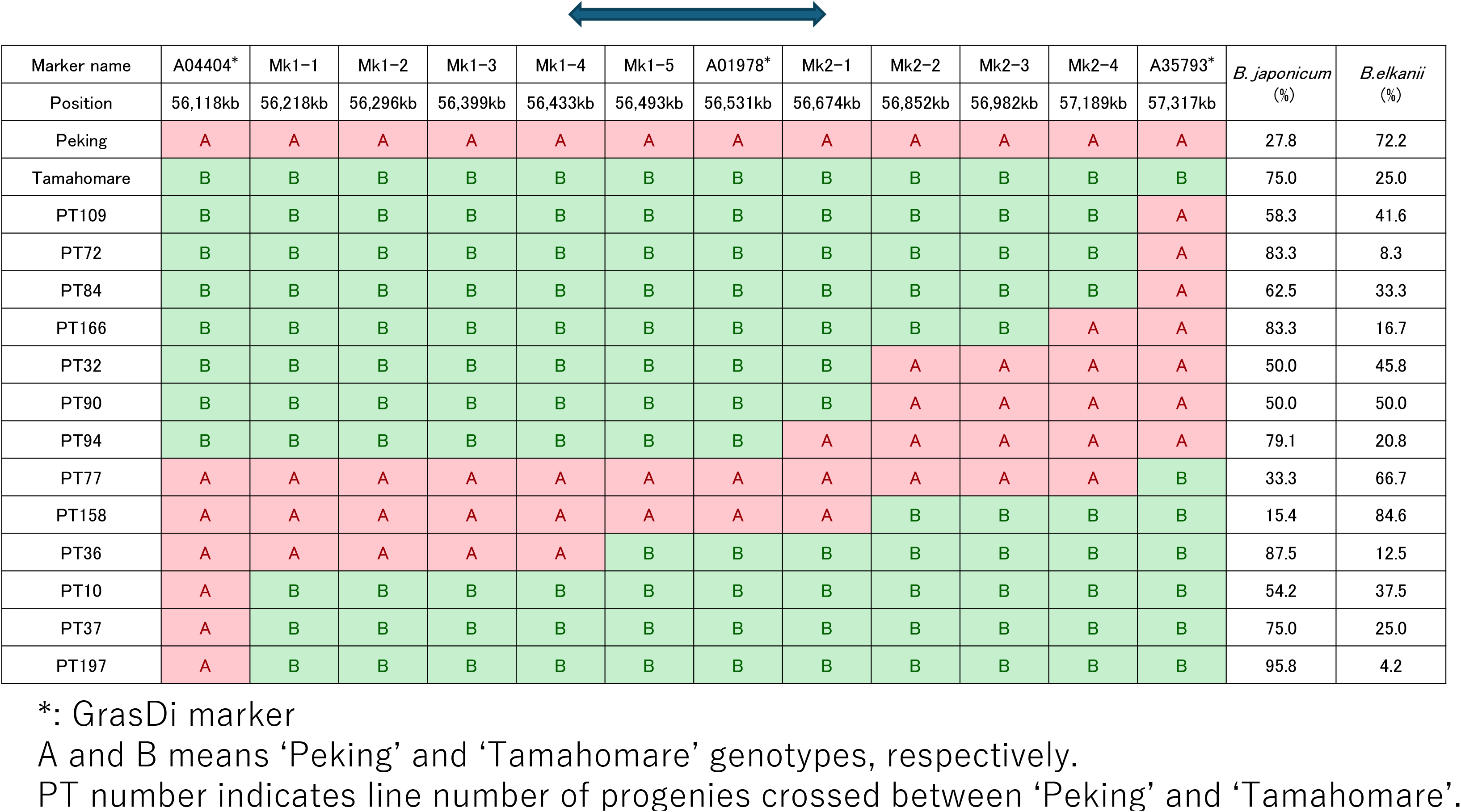
Fine mapping of the quantitative trait locus (QTL) on chromosome 18. *: GrasDi marker A and B means ‘Peking’ and ‘Tamahomare’ genotypes, respectively. PT number indicates line number of progenies crossed between ‘Peking’ and ‘Tamahomare’.

### 3.3 RNA-seq analysis and candidate genes

The RNA-seq analysis was performed using RNA derived from the roots of ‘Peking’ and ‘Tamahomare’ a month after germination to identify the potential candidate genes responsible for QTL. The results of the RNA-seq analysis revealed that only a single gene, Glyma.18g281700, was differentially expressed between the parents within the QTL region with padj<0.05 and fold change >2 (Figure S4; Table S3). The expression levels of Glyma.18g281700 were significantly higher in ‘Peking’ than in ‘Tamahomare.’ Glyma.18g281700 encodes NBS–ARC (apoptosis, R proteins, and CED-4) and LRR domains, often observed in disease resistance (R) genes.

## 4 DISCUSSION

Several genes regulating nodulation have been reported in soybeans. These are called *Rj*(s) and *rj*(s) genes, and *Rj2, Rj3, Rj4*, and *Rfg1* are the dominant alleles that restrict nodulation in specific strains (Hayashi et al., 2012), of which *Rj2* and *Rfg1* are allelic genes located on chromosome 16 and encode a member of the Toll/ interleukin-1 receptor–NBS–LRR resistance protein (Fan et al., 2017). The *Rj4* allele is located on chromosome 1 and encodes a thaumatin-like protein (Hayashi et al., 2014; Tang et al., 2014; Tang et al., 2016), and *Rj3* is restricted to nodulation by *B. elkanii* USDA33 (Vest 1970) but has not yet been identified. Other recessive genes that regulate nodulation induce nonnodulation (*rj1, rj5*, and *rj6*) (Pracht et al., 1993; Lee et al., 2011; Indrasumunar et al., 2011) and hypernodulation phenotypes (*rj7*) (Nishimura et al., 2002a; Nishimura et al., 2002b).

Ramongolalaina et al. (2018) demonstrated, for the first time, the chromosomal location of genes responsible for the proportion of nodulating bacterial strains in roots under natural non-inoculated conditions. In this study, we successfully narrowed down the location of the gene underlying the QTL to a 240 kb interval, wherein 22 predicted gene models were expected. RNA-seq revealed a single significantly differentially expressed gene between ‘Peking’ and ‘Tamahomare’ in the QTL interval.

Three consecutive genes, i.e., Glyma.18g281500, Glyma.18g281600, and Glyma.18g281700, have ubiquitin-like protease 1, NBS–ARC, and LRR domains, respectively (Jebanathirajah et al., 2002; Steele et al., 2019; Wei et al., 2023). These three tandemly arranged genes showed more than 90% sequence similarity. Glyma.18g281500 was located just outside the QTL region, and Glyma.18g281600 was within the QTL region, but these two genes did not exhibit a differential expression between the parents. Glyma.18g281700 alone differed substantially in its expression between the parents.

The genes *Rpp1* and *Rpp1b* that confer resistance to *Phakopsora pachyrhizi*, the causative agent of rust, are located within the same region on chromosome 18 (Pedley et al., 2019; Barros et al., 2023; Wei et al., 2023; Yamanaka et al., 2023). Pedley et al. (2019) reported that eight genes were homologous to the NBS–LRR family of disease R genes in four bacterial artificial chromosome contigs on chromosome 18, including the *Rpp1* locus. Three genes, namely Glyma.18g281500, 281600, and 281700, were considered he candidate genes for *Rpp1*, and Glyma.18g281600 was the most highly expressed candidate gene in PI200492. However, Barros et al. (2023) showed that the *Rpp1* gene in PI594756 was located immediately upstream of *Rpp1* in PI200492. Barros et al. (2023) indicated that genomic variations, such as presence/ absence and copy number variations, cause the diversification of disease R genes.

In conclusion, a single major QTL located on chromosome 18 was associated with the symbiotic compatibility of nodules. No genes implicated in symbiosis between rhizobia and soybeans were observed on chromosome 18. A single candidate gene was identified with differential transcription levels between ‘Peking’ and ‘Tamahomare.’ This candidate gene bears the NBS_–_LRR domain and has been reported as a candidate gene for soybean rust resistance, *Rpp1*.

## Supplemental data files

### Supplemental table titles

Table S1: Genetic map summary generated by GRAS-Di

Table S2: List of primer sequences used for fine mapping of the QTL located on Chromosome 18

Table S3: Summary of differential expressed genes analysis within the QTL region.

Table S4: Primers used for amplicon sequence

## Supplemental figure titles

Figure S1: QTL location on Chromosome 18

Figure S2: Distribution of the RILs for ratio of rhizobia strains used homozygous genotypes at the QTL region from the recombinant inbred lines from ‘Peking’ and ‘Tamahomare’

Figure S3: S3 Schematic view of position of molecular markers around the QTL region on Chromosome 18

Figure S4: `Heatmap of the 22 candidate genes in ‘Peking’ and ‘Tamahomare’`.

Figure S5: `Schematic representation of PCR amplification for ITS region`.

## ACKNOWLEDGMENTS

This work was supported by `a Grant-in-Aid for Scientific Research (C) (General) (for MT) (JSPS KAKENHI Grant number 22K05571) and` The Kyoto University Foundation (for MT).

We would like to thank Editage (www.editage.jp) for English language editing.

## AUTHOR CONTRIBUTIONS

MT conceived and designed the study. TY and TT developed the mapping population, constructed the genetic linkage map, and performed the fine mapping. TT and KS collected the phenotypic data. MT performed RNA-seq analysis and wrote the manuscript. All the authors contributed to the manuscript and approved the submitted version.

## DATA AVAILABILITY STATEMENT

The RNA-seq raw data used in this study were downloaded from the DRA database in the DDBJ under the BioProject accession number PRJDB17790.

