## supplementa file for "Identification of novel candidate genes associated with the symbiotic compatibility of soybean with rhizobia under natural conditions"

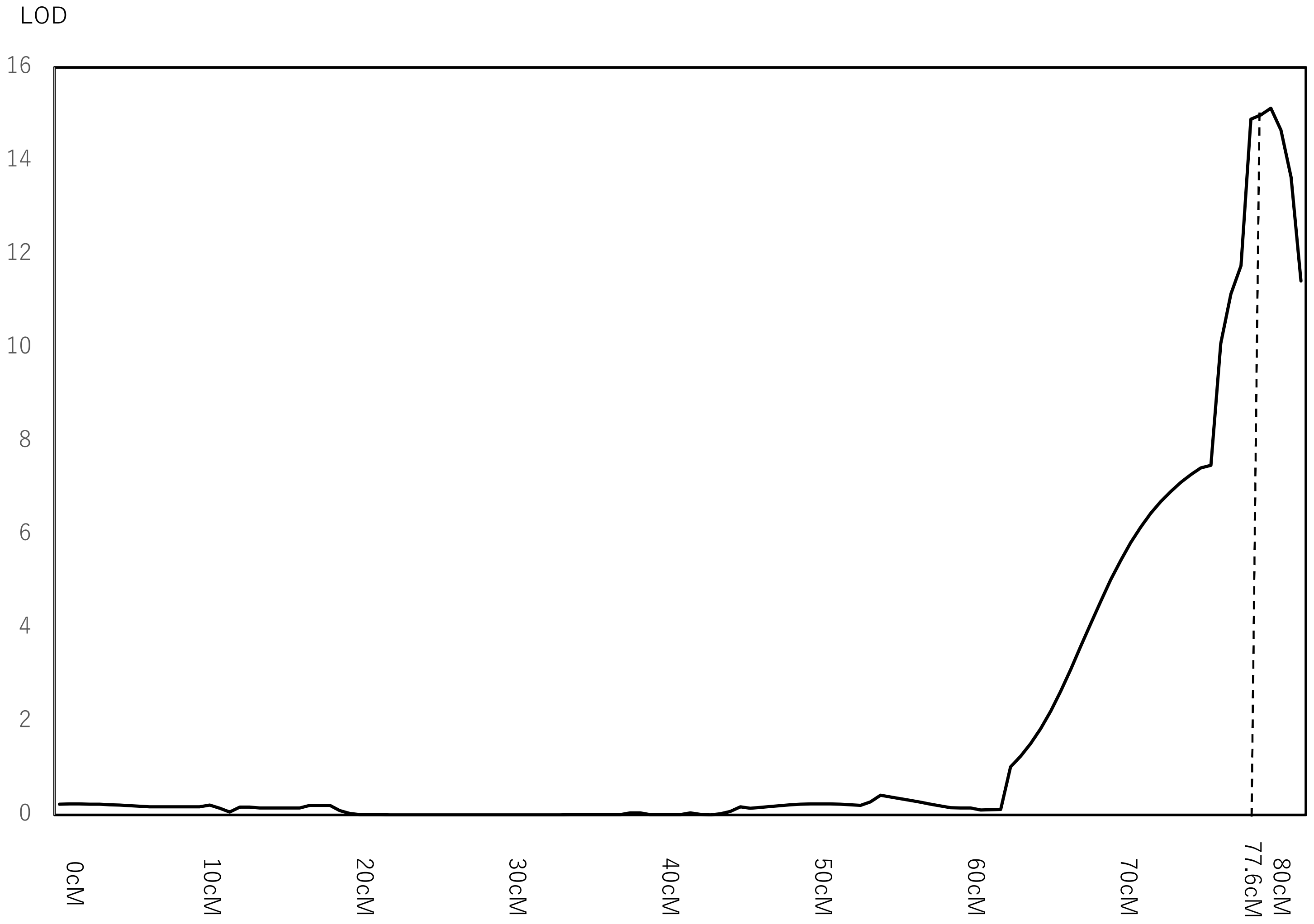

Figure S1  QTL location on Chromosome 18

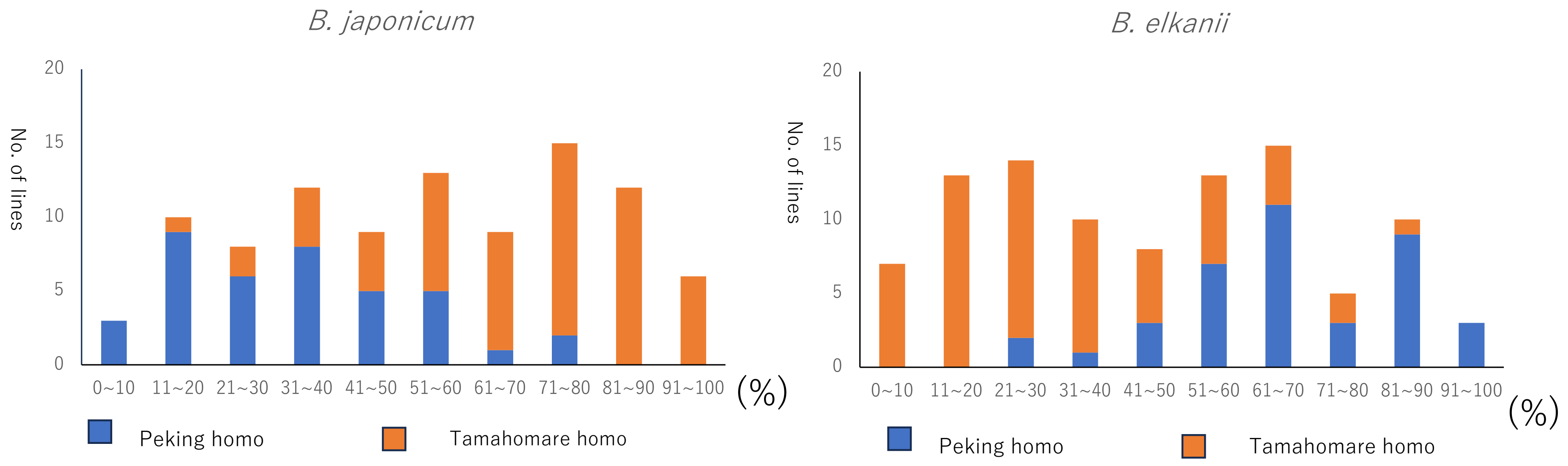

Figure S2 Distribution of the RILs for ratio of rhizobia strains used homozygous genotypes at the QTL region from the recombinant inbred lines from ‘Peking’ and ‘Tamahomare’

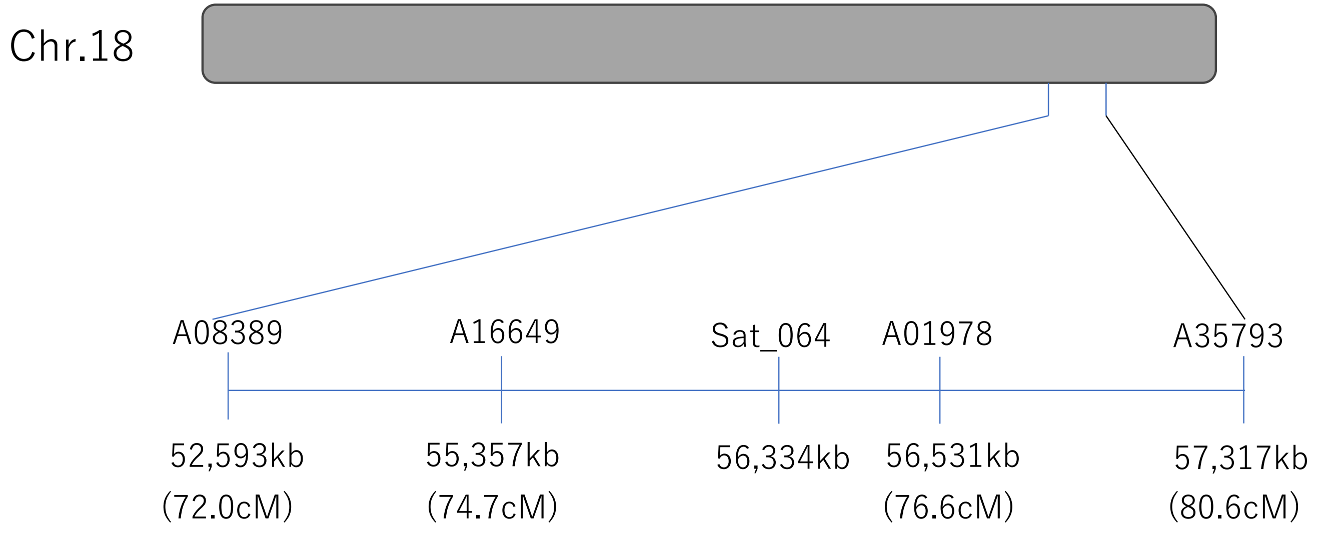

Figure S3 Schematic view of position of molecular markers around the QTL region on Chromosome 18

GrasDi marker: A08389, A16649, A01978, A35793
'Sat_064' is SSR marker used in Ramongolalaina et al (2018)

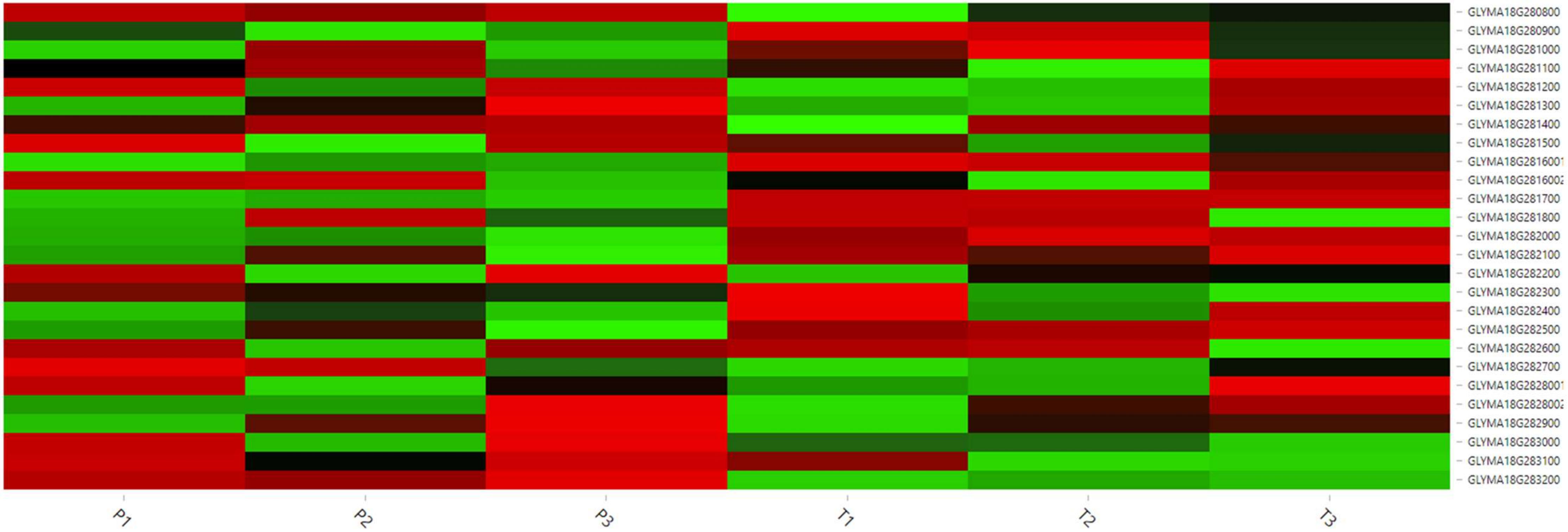

Figure S4 Heatmap of the 22 candidate genes in ‘Peking’ and ‘Tamahomare’.
The red blocks represent over expressed genes, and the green blocks represent under expressed genes.
P1-P3 indicated replicates of ‘Peking’, and T1-T3 indicated those of ‘Tamahomare’

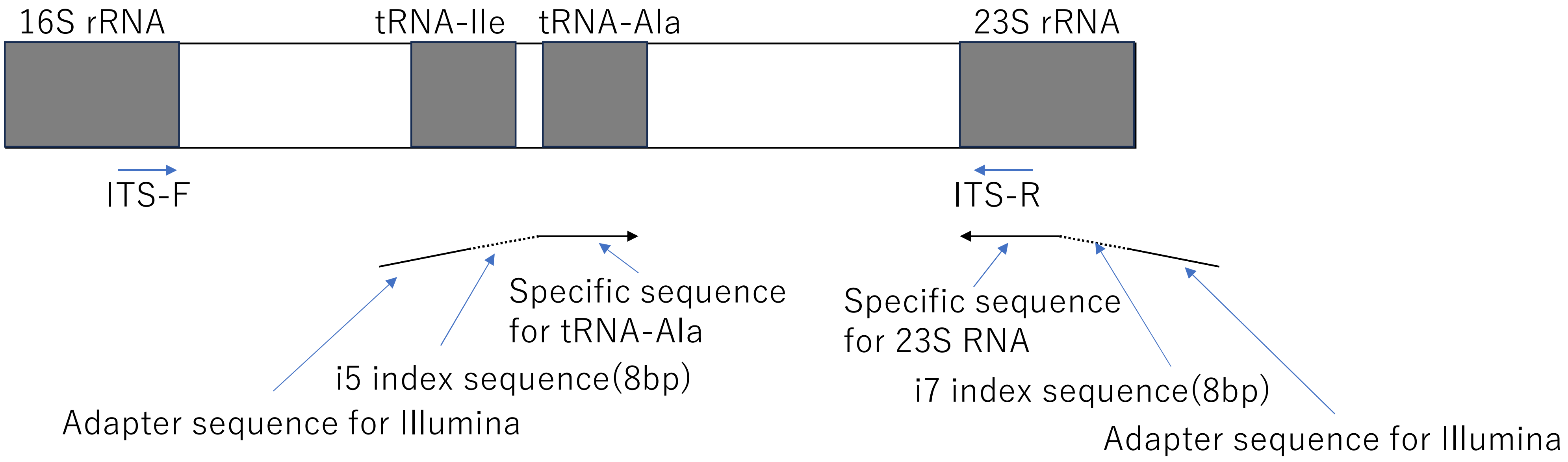

Figure S5 Schematic representation of PCR amplification for ITS region.
ITS region, amplified with ITS-F and ITS-R primers, is suitable for PCR-RFLP and sanger sequence. But amplified ITS region is app.900bp long, not suitable for shotgun amplicon sequence. Therefore, primer was designed in tRNA-Ala site.

### Table S1 Genetic map summary generated by GRAS-Di

| Chromosome No. | No. of marker | Total length（cM） | Average marker interval(cM) |
|---|---|---|---|
| 1 | 57 | 160.25 | 2.81 |
| 2 | 28 | 243.43 | 8.69 |
| 3 | 17 | 114.28 | 6.72 |
| 4 | 47 | 126.76 | 2.70 |
| 5 | 22 | 119.14 | 5.42 |
| 6 | 70 | 194.34 | 2.78 |
| 7 | 36 | 123.94 | 3.44 |
| 8 | 28 | 130.67 | 4.67 |
| 9 | 73 | 132.71 | 1.82 |
| 10 | 88 | 152.60 | 1.73 |
| 11 | 32 | 169.05 | 5.28 |
| 12 | 32 | 134.93 | 4.22 |
| 13 | 44 | 172.38 | 3.92 |
| 14 | 10 | 112.92 | 11.29 |
| 15 | 27 | 101.91 | 3.77 |
| 16 | 60 | 144.52 | 2.41 |
| 17 | 58 | 107.10 | 1.85 |
| 18 | 62 | 80.70 | 1.30 |
| 19 | 48 | 120.81 | 2.52 |
| 20 | 21 | 203.83 | 9.71 |
| ＊ | 860 | 2846.28 | 3.31 |

Table S2 List of primer sequences used for fine mapping of the QTL located on Chromosome 18

| Primer name | Position | Forward sequence | Reverse sequence |
|---|---|---|---|
| M1-1 | 56,218kb | CACTTGTGGTGCAATTCTTATCA | GGATATTCCCTGTTGTGTTGC |
| M1-2 | 56,296kb | CACCACACAGAGAAGCACAAA | GAGAGCCCAAGGCCATATAA |
| M1-3 | 56,399kb | AACCAGGGGTTTCTTTCTTTT | GACGGGTTGGATGTTATTGTG |
| M1-4(EcoRI) | 56,433kb | ATGCGTTGGGTTTAGGGTTA | CCTGCATCTCCTCCAAGTAAA |
| M1-5 | 56,493kb | AGATCATTGGATAACATCTGACTTT | CCATCTCAACCTCCTCCAGA |
| M2-1 | 56,674kb | ATCCCTCCTCTCTCCCTCTC | AAAATGCATTGTGGGCACTT |
| M2-2 | 56,852kb | CCTTCTCAAGTTAGCATAACAAAATG | GCTTCTATTGTGTCACTAACTGACTG |
| M2-3(EcoRV) | 56,982kb | CAATTCAATCGCTATTCGCTAGT | CACCCCAAAGTACCAGTTGC |
| M2-4(DraI) | 57,189kb | TTGACCGTGGCATATCTTCA | TGCGCCTAAAGATGCACTAA |

M1-4, M2-3 and M2-4 are Cleaved Amplified Polymorphic Sequence (CAPS) marker.
Other markers are indel marker.

### Table S3 Summary of differential expressed genes analysis within the QTL region.

| Gene model name[(1)] | Symbol[(2)] | Peking[(3)] | Tamahomare[(3)] | log2FC[(4)] | Padj[(5)] |
|---|---|---|---|---|---|
| GLYMA_18G280800 | KRH01498 | 0.032 | 0.336 | 0.373 | 0.495 |
| GLYMA_18G280900 | KRH01499 | 681.945 | 382.325 | -0.833 | 0.195 |
| GLYMA_18G281000 | KRH01500 | 2.060 | 1.276 | -0.427 | 0.637 |
| GLYMA_18G281100 | KRH01501 | 27.023 | 27.531 | 0.026 | 0.939 |
| GLYMA_18G281200 | KRH01502 | 127.081 | 161.024 | 0.339 | 0.517 |
| GLYMA_18G281300 | KRH01503 | 10.195 | 11.263 | 0.132 | 0.838 |
| GLYMA_18G281400 | KRH01504 | 1.258 | 5.777 | 1.586 | 0.215 |
| GLYMA_18G281500 | KRH01505 | 2.618 | 2.792 | 0.068 | 0.927 |
| GLYMA_18G2816001 | KRH01507 | 19.473 | 7.003 | -1.355 | 0.522 |
| GLYMA_18G2816002 | KRH01506 | 27.821 | 32.192 | 0.204 | 0.780 |
| GLYMA_18G281700 | KRH01508 | 4.887 | 0.208 | -2.286 | 0.000 |
| GLYMA_18G281800 | KRH01509 | 4.849 | 4.646 | -0.051 | 0.998 |
| GLYMA_18G282000 | KRH01512 | 11.368 | 7.062 | -0.617 | 0.080 |
| GLYMA_18G282100 | KRH01513 | 4.572 | 2.530 | -0.659 | 0.358 |
| GLYMA_18G282200 | KRH01514 | 4.061 | 4.658 | 0.161 | 0.837 |
| GLYMA_18G282300 | KRH01516 | 5.632 | 5.728 | 0.021 | 0.923 |
| GLYMA_18G282400 | KRH01520 | 8.551 | 7.535 | -0.162 | 0.933 |
| GLYMA_18G282500 | KRH01521 | 5.395 | 2.864 | -0.727 | 0.212 |
| GLYMA_18G282600 | KRH01522 | 4.179 | 4.298 | 0.033 | 0.973 |
| GLYMA_18G282700 | KRH01526 | 14.450 | 17.166 | 0.234 | 0.539 |
| GLYMA_18G2828001 | KRH01530 | 25.270 | 24.882 | -0.021 | 0.954 |
| GLYMA_18G2828002 | KRH01529 | 3.943 | 4.760 | 0.221 | 0.899 |
| GLYMA_18G282900 | KRH01531 | 0.727 | 0.897 | 0.136 | 0.895 |
| GLYMA_18G283000 | KRH01532 | 0.369 | 0.751 | 0.356 | 0.658 |
| GLYMA_18G283100 | KRH01533 | 13.127 | 16.080 | 0.274 | 0.476 |

(1) Gene model name referred to soybase (https://www.soybase.org)
(2) GenBank ID (3)average value of 3 replicate (4) Log 2 fold change (5) adjusted p-value

### Table S4 Primers  used for amplicon sequence

| primer name | sequence | length(bp) |
|---|---|---|
| IDT8_i5_01_RMOD_L | ACACTCTTTCCCTACACGACGCTCTTCCGATCTATATGCGCGTCTGGCTCCACCAGATG | 59 |
| IDT8_i5_02_RMOD_L | ACACTCTTTCCCTACACGACGCTCTTCCGATCTTGGTACAGGTCTGGCTCCACCAGATG | 59 |
| IDT8_i5_03_RMOD_L | ACACTCTTTCCCTACACGACGCTCTTCCGATCTAACCGTTCGTCTGGCTCCACCAGATG | 59 |
| IDT8_i5_04_RMOD_L | ACACTCTTTCCCTACACGACGCTCTTCCGATCTTAACCGGTGTCTGGCTCCACCAGATG | 59 |
| IDT8_i5_05_RMOD_L | ACACTCTTTCCCTACACGACGCTCTTCCGATCTGAACATCGGTCTGGCTCCACCAGATG | 59 |
| IDT8_i5_06_RMOD_L | ACACTCTTTCCCTACACGACGCTCTTCCGATCTCCTTGTAGGTCTGGCTCCACCAGATG | 59 |
| IDT8_i5_07_RMOD_L | ACACTCTTTCCCTACACGACGCTCTTCCGATCTTCAGGCTTGTCTGGCTCCACCAGATG | 59 |
| IDT8_i5_08_RMOD_L | ACACTCTTTCCCTACACGACGCTCTTCCGATCTGTTCTCGTGTCTGGCTCCACCAGATG | 59 |
| IDT8_i5_09_RMOD_L | ACACTCTTTCCCTACACGACGCTCTTCCGATCTAGAACGAGGTCTGGCTCCACCAGATG | 59 |
| IDT8_i5_10_RMOD_L | ACACTCTTTCCCTACACGACGCTCTTCCGATCTTGCTTCCAGTCTGGCTCCACCAGATG | 59 |
| IDT8_i5_11_RMOD_L | ACACTCTTTCCCTACACGACGCTCTTCCGATCTCTTCGACTGTCTGGCTCCACCAGATG | 59 |
| IDT8_i5_12_RMOD_L | ACACTCTTTCCCTACACGACGCTCTTCCGATCTCACCTGTTGTCTGGCTCCACCAGATG | 59 |
| IDT8_i5_13_RMOD_L | ACACTCTTTCCCTACACGACGCTCTTCCGATCTATCACACGGTCTGGCTCCACCAGATG | 59 |
| IDT8_i5_14_RMOD_L | ACACTCTTTCCCTACACGACGCTCTTCCGATCTCCGTAAGAGTCTGGCTCCACCAGATG | 59 |
| IDT8_i5_15_RMOD_L | ACACTCTTTCCCTACACGACGCTCTTCCGATCTTACGCCTTGTCTGGCTCCACCAGATG | 59 |
| IDT8_i5_16_RMOD_L | ACACTCTTTCCCTACACGACGCTCTTCCGATCTCGACGTTAGTCTGGCTCCACCAGATG | 59 |
| IDT8_i5_17_RMOD_L | ACACTCTTTCCCTACACGACGCTCTTCCGATCTATGCACGAGTCTGGCTCCACCAGATG | 59 |
| IDT8_i5_18_RMOD_L | ACACTCTTTCCCTACACGACGCTCTTCCGATCTCCTGATTGGTCTGGCTCCACCAGATG | 59 |
| IDT8_i5_19_RMOD_L | ACACTCTTTCCCTACACGACGCTCTTCCGATCTGTAGGAGTGTCTGGCTCCACCAGATG | 59 |
| IDT8_i5_20_RMOD_L | ACACTCTTTCCCTACACGACGCTCTTCCGATCTACTAGGAGGTCTGGCTCCACCAGATG | 59 |
| IDT8_i5_21_RMOD_L | ACACTCTTTCCCTACACGACGCTCTTCCGATCTCACTAGCTGTCTGGCTCCACCAGATG | 59 |
| IDT8_i5_22_RMOD_L | ACACTCTTTCCCTACACGACGCTCTTCCGATCTACGACTTGGTCTGGCTCCACCAGATG | 59 |
| IDT8_i5_23_RMOD_L | ACACTCTTTCCCTACACGACGCTCTTCCGATCTCGTGTGTAGTCTGGCTCCACCAGATG | 59 |
| IDT8_i5_24_RMOD_L | ACACTCTTTCCCTACACGACGCTCTTCCGATCTGTTGACCTGTCTGGCTCCACCAGATG | 59 |
| IDT8_i7_01_RMOD_R | GTGACTGGAGTTCAGACGTGTGCTCTTCCGATCTCTGATCGTCCTTCATCGCCTCTCA | 58 |
| IDT8_i7_02_RMOD_R | GTGACTGGAGTTCAGACGTGTGCTCTTCCGATCTACTCTCGACCTTCATCGCCTCTCA | 58 |
| IDT8_i7_03_RMOD_R | GTGACTGGAGTTCAGACGTGTGCTCTTCCGATCTTGAGCTAGCCTTCATCGCCTCTCA | 58 |
| IDT8_i7_04_RMOD_R | GTGACTGGAGTTCAGACGTGTGCTCTTCCGATCTGAGACGATCCTTCATCGCCTCTCA | 58 |
| IDT8_i7_05_RMOD_R | GTGACTGGAGTTCAGACGTGTGCTCTTCCGATCTCTTGTCGACCTTCATCGCCTCTCA | 58 |
| IDT8_i7_06_RMOD_R | GTGACTGGAGTTCAGACGTGTGCTCTTCCGATCTTTCCAAGGCCTTCATCGCCTCTCA | 58 |
| IDT8_i7_07_RMOD_R | GTGACTGGAGTTCAGACGTGTGCTCTTCCGATCTCGCATGATCCTTCATCGCCTCTCA | 58 |
| IDT8_i7_08_RMOD_R | GTGACTGGAGTTCAGACGTGTGCTCTTCCGATCTACGGAACACCTTCATCGCCTCTCA | 58 |
| IDT8_i7_09_RMOD_R | GTGACTGGAGTTCAGACGTGTGCTCTTCCGATCTCGGCTAATCCTTCATCGCCTCTCA | 58 |
| IDT8_i7_10_RMOD_R | GTGACTGGAGTTCAGACGTGTGCTCTTCCGATCTATCGATCGCCTTCATCGCCTCTCA | 58 |
| IDT8_i7_11_RMOD_R | GTGACTGGAGTTCAGACGTGTGCTCTTCCGATCTGCAAGATCCCTTCATCGCCTCTCA | 58 |
| IDT8_i7_12_RMOD_R | GTGACTGGAGTTCAGACGTGTGCTCTTCCGATCTGCTATCCTCCTTCATCGCCTCTCA | 58 |
| IDT8_i7_13_RMOD_R | GTGACTGGAGTTCAGACGTGTGCTCTTCCGATCTTACGCTACCCTTCATCGCCTCTCA | 58 |
| IDT8_i7_14_RMOD_R | GTGACTGGAGTTCAGACGTGTGCTCTTCCGATCTTGGACTCTCCTTCATCGCCTCTCA | 58 |
| IDT8_i7_15_RMOD_R | GTGACTGGAGTTCAGACGTGTGCTCTTCCGATCTAGAGTAGCCCTTCATCGCCTCTCA | 58 |
| IDT8_i7_16_RMOD_R | GTGACTGGAGTTCAGACGTGTGCTCTTCCGATCTATCCAGAGCCTTCATCGCCTCTCA | 58 |
| IDT8_i7_17_RMOD_R | GTGACTGGAGTTCAGACGTGTGCTCTTCCGATCTGACGATCTCCTTCATCGCCTCTCA | 58 |
| IDT8_i7_18_RMOD_R | GTGACTGGAGTTCAGACGTGTGCTCTTCCGATCTAACTGAGCCCTTCATCGCCTCTCA | 58 |
| IDT8_i7_19_RMOD_R | GTGACTGGAGTTCAGACGTGTGCTCTTCCGATCTCTTAGGACCCTTCATCGCCTCTCA | 58 |
| IDT8_i7_20_RMOD_R | GTGACTGGAGTTCAGACGTGTGCTCTTCCGATCTGTGCCATACCTTCATCGCCTCTCA | 58 |
| IDT8_i7_21_RMOD_R | GTGACTGGAGTTCAGACGTGTGCTCTTCCGATCTGAATCCGACCTTCATCGCCTCTCA | 58 |
| IDT8_i7_22_RMOD_R | GTGACTGGAGTTCAGACGTGTGCTCTTCCGATCTTCGCTGTTCCTTCATCGCCTCTCA | 58 |
| IDT8_i7_23_RMOD_R | GTGACTGGAGTTCAGACGTGTGCTCTTCCGATCTTTCGTTGGCCTTCATCGCCTCTCA | 58 |
| IDT8_i7_24_RMOD_R | GTGACTGGAGTTCAGACGTGTGCTCTTCCGATCTAAGCACTGCCTTCATCGCCTCTCA | 58 |
